# Generalizable Neural Models of Emotional Engagement and Disengagement

**DOI:** 10.1101/2024.02.15.579332

**Authors:** Melanni Nanni-Zepeda, Travis C. Evans, Audreyana Jagger-Rickels, Gal Raz, Talma Hendler, Yan Fan, Simone Grimm, Martin Walter, Michael Esterman, Agnieszka Zuberer

**Author notes:** Author Note Correspondence concerning this article should be addressed to Melanni Nanni-Zepeda, Department of Psychiatry and Psychotherapy, Attention and Affect Laboratory, Eberhard Karls University, Tübingen, Germany. and.

## Abstract

Emotional experiences are never static but continuously evolve in response to internal and external contexts. Little is known about how neural patterns change as a function of these experiences, particularly in response to complex, real-world stimuli. This study aimed to identify generalizable neural patterns as individuals collectively engage and disengage from emotions dynamically. To do so, we analyzed functional magnetic resonance imaging (fMRI) along with subjective emotional annotations from two independent studies as individuals watched negative and neutral movie clips. We used predictive modeling to test if a model trained to predict a group emotional signature response in one study generalizes to the other study and vice versa. Disengagement patterns generalized specifically across intense clips. They were supported by connections within and between the sensorimotor and salience networks, maybe reflecting the processing of feeling states as individuals regulate their emotions. Prediction success for the engagement signature was mixed, but primarily linked to connections within the visual and between the visual and dorsal attention networks, maybe supporting visual attention orienting as emotions intensify. This work offers potential pathways for identifying generalizable neural patterns contributing to future affective research and clinical applications aiming to better understand dynamic emotional responses to naturalistic stimuli.

## Introduction

Our well-being profoundly depends on how we engage *in* and disengage *from* emotions when interacting with others and the world. While theories of affect emphasize the temporal progression of emotional responses and their regulation (e.g. Gross et al. (2014); Scherer (2009); Sheppes and Gross (2011) pp. 16-17), these processes have typically been studied in isolation (Goldin et al., 2009; Gruber et al., 2011; McRae et al., 2010), disregarding their dynamic nature throughout time. In real life, emotions evolve, and naturalistic scenarios like movies may preserve this temporal alternation of engagement and disengagement, offering rich, dynamic affective experiences (Morgenroth et al., 2023; Saarimäki, 2021). Although movies have the potential to mimic real-life scenarios, there is a challenge for affective movie fMRI in terms of ecological validity not necessarily directly translating to ecological generalizability, in that affective correlates may not extend uniformly across various naturalistic contexts (Nastase et al., 2020), and instead may be partially driven by contextual factors (jump cuts, actor’s costumes, camera perspective, background) (Hasson et al., 2008; Zacks & Magliano, 2011). Predictive modeling of a dynamic affective experience across various movies and participant groups holds the potential to unveil more generalized neural mechanisms relevant to the actual emotional experience, independent from the idiosyncratic nature of each movie. This way, common patterns, and neural responses can be distilled that transcend specific cinematic content and individual differences. As stimuli continuously evolve, individuals dynamically allocate attentional resources toward affectively relevant information, engaging with and eventually disengaging from specific content. Theoretical work suggests that emotional intensity can be understood through both trajectories of intensity over time (gradual changes) and episodes of sharp increases and decreases of intensity (Kuppens et al., 2010). Building on this framework, we propose that episodes can be conceptualized as containing two distinct components: an *engagement component*, marked by the sharp increase in emotional intensity, followed by a *disengagement component*, characterized by the subsequent decrease in intensity. These cyclical engagement-disengagement patterns would together contribute to the formation of subjective emotional experiences. Specifically, arousal would function as the energetic substrate that enables the formation of top-down attentional biases toward emotionally salient stimuli (Mohanty et al., 2008), amplifying competitive processing advantages for high-priority information while suppressing lower-priority competing representations (Mather & Sutherland, 2011). In this context, emotional engagement would not reflect simply ’more arousal’, but instead reflect a transitional process of directing and intensifying attention towards movie content. In contrast, disengagement would reflect the transition from high to low arousal as the arousal-mediated competitive bias dissolves, allowing attention to withdraw from movie content and return to a more distributed, less focused processing state. In other words, two moments in time may share the same arousal level yet reflect different cognitive processes, one reflecting passive arousal without competitive advantage, and the other reflecting arousal actively deployed to bias processing in favor of emotionally salient movie content. Together, continuous intensity tracking techniques may provide a unique opportunity to identify moments of engagement and disengagement within the stream of emotional experiences and detect underlying generalizable brain patterns. Few studies have shown that continuous subjective reports of attention and emotion state can be predicted across movies and participant groups. One study showed that continuous subjective experience of fear during movie viewing was predicted by widespread large-scale brain network connectivity patterns, and that these predictions were successful between independent subject groups and movies (Zhou et al., 2023). In one study it was found that functional connectivity strength within the DMN was predictive of continuous attentional engagement reports across a movie and audiobook from independent studies (Song, Finn, et al., 2021). Other work linked DMN connectivity to both intensity and polarity of continuous affective experiences across independent movie data sets (Lettieri et al., 2022). One study found that even though stronger dynamic DMN recruitment was also linked to moment-to-moment movie comprehension in separate movies, a DMN signature obtained in one movie failed to predict movie comprehension in the other movie and vice versa (Song, Park, et al., 2021). Together, these findings suggest that both dynamic changes in attentional engagement levels and affective processing, or an interplay of both may be tracked by dynamic functional brain connectivity (dynFC) across large-scale networks. While this body of work demonstrates that continuous annotations can predict generalizable brain patterns across movies, those underlying collective engagement and disengagement as distinct episodic processes defined by transitional intensity changes rather than absolute levels remain poorly understood.

To tackle this, we analyzed movie fMRI data from two independent studies, both using retroactive subjective emotional intensity annotations in response to either one intense negative and one neutral movie clip (study 1) or one intense negative movie clip (study 2). Positive and negative shifts in subjects’ annotation timecourses were labeled as EE and ED, respectively. We used support vector regression (SVR) to test the predictive nature of dynFC patterns across independent data linked to moment-to-moment changes in the degree to which a participant group exhibited EE and ED phases throughout movie viewing. Based on previous work linking subjective attention states during movie viewing to the DMN, we wondered if DMN connectivity, both within and between networks, might play a central role in predicting group signature of EE or ED. However, given the limited predictive studies in naturalistic scenarios, we did not constrain our analysis to specific regions or networks of interest.

We hypothesized that collective emotional engagement (EE) and emotional disengagement (ED) during intense negative movies are linked to brain patterns generalizable across independent participant groups and movie clips but not generalizable to neutral movie content of low emotional intensity, suggesting high intensity-specific predictive brain patterns. Our results were in line with this for predicting ED signatures, where predictions succeeded between negative movies and failed for between negative and neutral predictions.

In contrast, our results were mixed for predicting EE signatures, where predictions succeeded between negative movies and also partially for between negative and neutral predictions. Brain patterns predicting collective disengagement involved functional connectivity patterns across the entire brain connectome without drawing on specific network-to-network connections. In contrast, EE signatures were linked to connections from visual to attention and salience networks, potentially reflecting heightened engagement.

This work represents an initial step toward understanding generalizable neural patterns underlying collective engagement and disengagement patterns during naturalistic viewing.

## Material and Methods

### Data Sources

We analyzed fMRI and behavioral data from two independent studies with similar paradigms and acquisition methods. These prior studies addressed unrelated research questions. For clarity, we refer to them as study 1 (Borchardt et al., 2018) and study 2 (Raz et al., 2016). We previously published work based on study 1 (Nanni-Zepeda et al., 2024) with another research question.

### Participants

Study 1 included 22 female volunteers aged 20–49 years (mean age 28. 1 ∓ 6.5), all German native speakers recruited in Germany. Study 2 included 44 Hebrew native speakers recruited in Tel Aviv, Israel (25 female, 19 male) aged 21–37 years (mean age 26.73 ∓ 4.69). All participants from both studies were screened for neurological and psychiatric disorders using standardized protocols (study 1: short SCID (Wittchen et al., 1997)), with no participants reporting any such conditions. Study 1 protocols were conducted in accordance with the Declaration of Helsinki and approved by the institutional review board of the Charité. Study 2 was reviewed and approved by the ethics committees of the Tel Aviv Sourasky Medical Center.

### Naturalistic Viewing Paradigm

Participants in study 1 watched a neutral and negative movie clip in a counterbalanced order during fMRI scanning. The negative clip (*21 Grams*, Iñárritu (2003), 4.45 min) depicts a mother learning of her daughters’ deaths, while the neutral clip (Son’s Room, Moretti (2001); 4.54 min) shows everyday family life, matched for low-level features such as faces and domestic settings. Both were dubbed in German. The negative clip elicited significantly stronger negative arousal and valence (Borchardt et al., 2018).

In study 2, participants watched a 10-min excerpt from Sophie’s Choice (Pakula, 1983), where a mother must choose which child to save, containing similar visual elements (faces, domestic context). It was presented in English with Hebrew subtitles. These stimuli have been validated in prior fMRI emotion-induction studies (Borchardt et al., 2018; Gaviria et al., 2021; Hanich et al., 2014; Innes-Ker, 2015; Raz et al., 2016).

### Subjective Continuous Emotional Intensity Annotations

About 15 minutes post-scan, they re-watched the clips while providing continuous emotional intensity annotations via a trackball-operated mouse. A visual analog scale (VAS) from 0 ("not at all") to 250 ("very much") was displayed alongside the video for real-time adjustments (instructions in supplements). Annotations were sampled at 30Hz and downsampled to the fMRI sampling rate. Study 2 followed a similar procedure.

Participants viewed the clip during fMRI scanning and re-watched it 15 minutes later, providing continuous emotional intensity annotations using a 7-point Likert scale (0 = "neutral" to 7 = "very intense"). For details, see Raz et al. (2016). Annotations were sampled at 10Hz and downsampled to match the fMRI sampling rate. In both studies, individuals were instructed to annotate their emotional intensity with regard to the first time they watched the movie. Figure 1 shows the variation in emotional intensity ratings over time across subjects.

**Figure 1.**
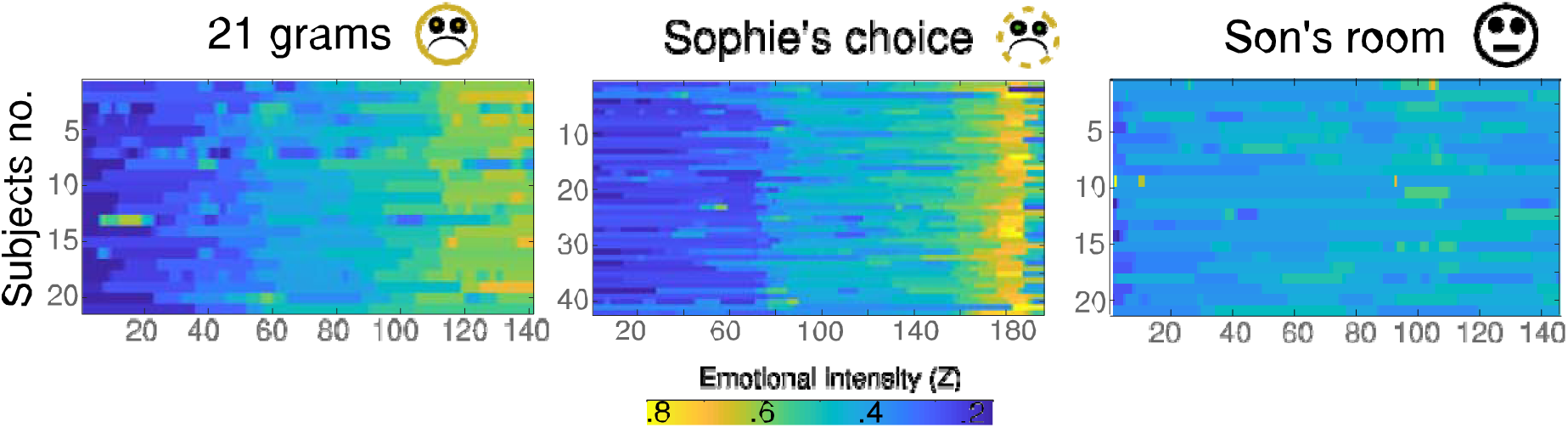
(A) Individual subjective emotional intensity annotations over time for each movie clip, z-scored and scaled between 0 and 1 for visualization purposes. *EE*, Emotional Engagement, *ED* Emotional Disengagement.

**Figure 2.**
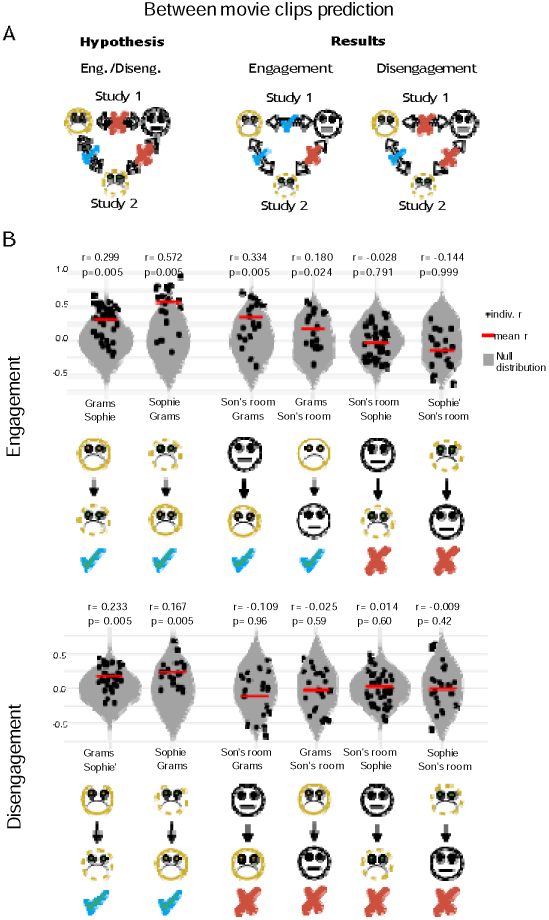
Functional connectivity patterns underlying engagement and disengagement generalize across independent studies (A) left panel: To infer specificity to negative clips, models trained to predict a group signature response of EE and ED in one negative movie clip in one study (e.g. study 1) must succeed in predicting the group signature response in another negative movie clip (e.g. study 2) and vice versa, but fail for predictions across negative and neutral clips. Based on these assumptions we infer generalization due to both independence from idiosyncratic features of the participant group and narratives in each study. Right panel: Both conditions were met for the group signature response of ED. For EE, only the first condition (between negative clip prediction) was met. (B) Predictive performance of dynFC features shown for each clip separately. The black dots in the scatter plots represent Pearson’s correlations between predicted and observed group signature response for EE (top) and ED (bottom). Horizontal lines indicate the prediction quality reflected in the mean r across cross-validation folds. Violin plots illustrate the null distributions of mean prediction performance after phase-randomization. The significance of empirical r was computed based on the null distribution (one-tailed t-test). *DynFC*, Dynamic Functional Connectivity, *EE*, Emotional Engagement, *ED*, Emotional Disengagement.

We used the individual subjective annotations to compute group-averaged signals of EE and ED through a sliding window approach, aggregating all instances within each sliding window. These group-averaged measures were used as input variables in our computational models, as detailed in the ’Emotional Engagement and Disengagement’ section.

### Emotional Engagement and Disengagement

We extracted time points of EE and ED from individual subjective annotations with an algorithm previously used to separate four phases of a temporal signal into rise, high, fall and low magnitude (Dessu et al., 2020; Kato et al., 2015; Shine et al., 2019). EE and ED periods were defined as rises and falls within the temporal fluctuations of subjective emotional intensity. To count as an EE or ED period, an emotional intensity score had to be within the 33rd and 67th percentile and exhibit a positive (EE) or a negative change (ED) between two subsequent time points. This way we obtained two binary time courses per participant: one indicating the presence of EE (yes/no), and the other indicating ED (yes/no) at each moment. Figure S1 shows the distribution of estimated EE and ED per movie.

We next sought to assess the degree to which individuals collectively exhibited EE and ED phases throughout movie viewing. To this aim, we obtained a dynamic group signature reflecting the dynamic change in frequency of simultaneous reports of EE or ED across individuals. We used a sliding window approach over the binary timeseries of EE and ED (window size of 40s; time step of 1 TR) where we counted the number of EE or ED moments present across all individuals within each sliding window. This resulted in two group signature time courses, i.e. one for EE, and one for ED. These time courses were then convolved with the hemodynamic response function (HRF) and used in combination with fMRI timecourses to train and test SVR models (Fig. 3).

**Figure 3.**
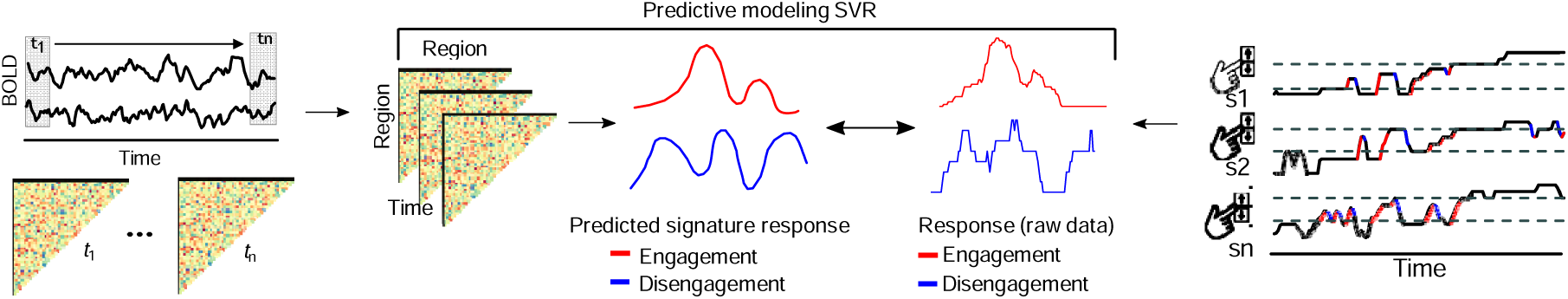
Visualization of how dynFC and group signature responses for EE and ED were derived. (A) DynFC and the group signature responses were calculated using a sliding window approach. From left to right: DynFC represents the Pearson correlation of BOLD time courses within each sliding window, while the group signature responses for EE and ED reflects the number of EE and ED phases within each sliding window aggregated across all participants. An EE and ED phase was identified as positive or negative shifts within a subject’s continuous subjective emotional intensity annotation (see methods section). *DynFC*, Dynamic Functional Connectivity, *EE*, Emotional Engagement, *ED*, Emotional Disengagement.

To assess the robustness of our engagement and disengagement predictions, we conducted a sensitivity analysis testing whether the results held across varying percentile thresholds used to define EE and ED. The original threshold range (33rd and 67th percentiles) was systematically modified to include both wider and narrower ranges. We expanded the range in three incremental steps by decreasing the low percentile and increasing the high percentile by 5-unit increments, resulting in threshold pairs of (28th, 72nd), (23rd, 77th), and (18th, 82nd percentiles). We also tested a narrower range (38th, 62nd percentiles). For each threshold pair, we repeated the complete prediction analysis pipeline and evaluated the consistency of our main findings. Results are presented in supplementary materials (Tables S1 and S2).

### fMRI Analysis

fMRI data were acquired on a 3T scanner and preprocessed using standard pipelines (see Supplementary for full details).

#### Dynamic Predictive Modeling

To extract dynFC, for every participant, we extracted BOLD time courses using a 200 region parcellation by Schaefer et al. (2018), the Harvard-Oxford structural subcortical atlas (8 regions) Kennedy et al. (2016) and insula sub-regions (6 regions) Deen et al. (2011) resulting in a total of 214 brain regions. Next, we calculated dynFC of BOLD time series, computed as the Pearson correlation between pairs of regions (214 x 214 ROIs). We used a tapered sliding window with a length of 40 seconds and a tapering factor of 1 TR as done previously in naturalistic studies linking dynFC to subjective annotations (Petrican et al., 2021; Song, Finn, et al., 2021; Song, Park, et al., 2021).

In the next step, we used support vector regression (SVR) to test whether dynamic inter-regional connectivity patterns could predict temporal fluctuations in the group-level EE and ED signatures.

To reduce dimensionality and enhance model performance, we first identified functional connections (ROI pairs) that significantly correlated with the group EE or ED signature (one-sample t-test, p < 0.01). Using these selected connections as features, we trained an SVR model with a radial basis function (RBF) kernel (maximum iterations = 1,000; sklearn.svm.SVR in Python) to predict the group engagement time series based on each participant’s individual dynamic functional connectivity (dynFC) time courses.

To examine the generalizability of the identified engagement-related connectivity features, we performed across-dataset analyses. Specifically, feature weights derived from the within-movie analyses in one dataset were applied to predict the group EE and ED signatures in a different dataset, and vice versa. This procedure was also repeated for neutral versus negative clips to test whether predictive connectivity patterns were specific to emotionally intense narratives (Fig. 2A). Statistical significance of the mean correlation values was assessed by comparing them to a null distribution derived from SVR models trained on phase-randomized EE and ED time series.

Predictive performance was assessed by computing the Pearson correlation between the predicted and actual group engagement time series for each participant. For comparison, we also computed mean squared error (MSE) and the coefficient of determination (R2), where the R2 corresponds to the standard coefficient of determination, which quantifies how much variance in the observed engagement time series is explained by the model. We computed R^2^ using the standard coefficient of determination:

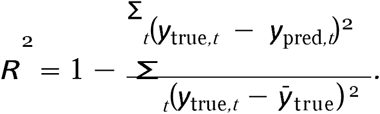

We averaged these metrics across participants to summarize group-level model accuracy. All analyses used publicly shared code from Song, Finn, et al. (2021) to ensure reproducibility (https://github.com/hyssong/NarrativeEngagement).

We also report within-dataset predictions, in which the group engagement signature was recalculated using a leave-one-out (LOO) approach. For each iteration, the SVR model was trained on all subjects except one, using the group signature computed without the held-out subject. The trained model was then applied to the held-out subject’s dynFC data to predict that same group-defined engagement time series (see Supplementary) To assess the significance of the empirical mean correlation values, we computed the percentage of simulated correlation values resulting from the SVR phase-randomized EE and ED. The threshold was set to ensure that this percentage did not surpass 95% (p < 0.05); Fig. 2B. This iterative procedure was repeated 200 times. Throughout this section’s analyses, we employed a modified version of the publicly available dynamic predicting modeling code initially developed by Song et al. (2021).

#### Networks Supporting Engagement and Disengagement

We further explored the involvement of canonical networks in predicting group signature response of EE and ED. We did this by grouping connections that significantly contributed to the model into canonical functional networks based on the network parcellation by Yeo et al. (2011). We then calculated the proportion of all participants’ significant connections against all possible connections within a particular network. To evaluate the relevance of each network in predicting a group signature response of EE and ED, we generated null matrices by phase-randomizing the empirical emotional response time courses. This process involved transforming the original data into the frequency domain, randomizing the phases, and then transforming the data back into the time domain to generate surrogate data with equivalent second-order properties as the original time series (Gias, 2023). These randomized data sets were input into the SVR model to generate simulated predictions, forming the null distribution. The significance of empirical mean r values was assessed by comparing the observed proportion of network pairs in the empirical data to the proportion of network pairs from the randomized data that surpassed the observed value. We inferred significance if more than 99% of the randomized pairs exceeded the empirical proportion (p < 0.01); Fig. 4A. We then grouped the significant network pairs into networks (Fig. 4B) and identified the region pairs predictive of EE and ED between movie clips in over 50% of the subjects. For visualization, we selected the two top nodes that exhibited most connections to other regions across the connectome (Fig. 4C).

**Figure 4.**
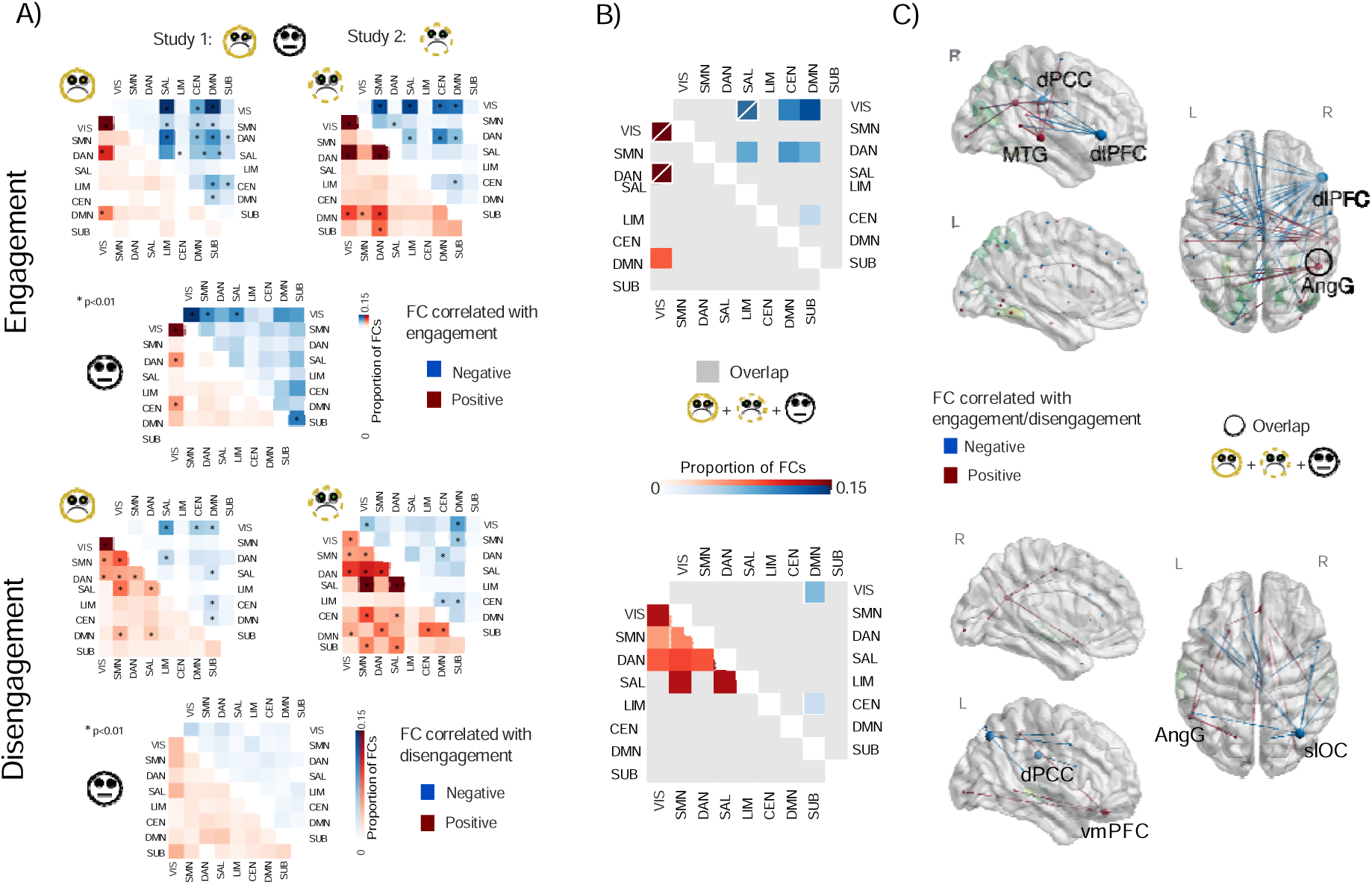
Functional connectivity within and across canonical networks predicts a group signature response of EE and ED. (A) predictive between-region connections were grouped into canonical functional networks (Schaefer et al., 2018). The lower triangle matrix (red) illustrates the proportion of FCs positively correlated with the group signature response for EE or ED, and the upper triangle matrix (blue) illustrates the proportion of FCs negatively correlated with the group signature response for EE or ED. Asterisks indicate network pairs selected above chance (one-tailed t tests, p_FDR_< 0.001). The intensity of colors indicates the proportion of selected connections from all possible ROI-to-ROI connections with regard to its network assignment. (B) Shared significant ROI-to-ROI connections across negative movie clips. The crossed gray square indicates connections shared among negative movie clips that also overlapped with the neutral clip. (C) Top two strongest ROI-to-ROI connections predicting a group signature response for EE and ED. Connections overlapping across all three movie clips are indicated by black circles, whereas those specific to negative clips do not show black circles. *EE*, Emotional Engagement, *ED*, Emotional Disengagement.

## Results

### Functional Connectivity Patterns underlying Engagement and Disengagement Generalize across Studies

In a next step, we investigated whether the neural features predicting the group signature response of EE and ED were generalizable across movie clips. To this aim, we tested if a model trained to predict a group signature response in one study is able to predict the group signature response in the other study and vice versa (Fig. 2). First, we tested predictions across negative clips of study 1 and 2. The model trained on dynFC of subjects watching the negative clip of study 2 successfully predicted the EE group signature in the negative clip of study 1 (r = 0.572; p < 0.005, MSE = 50.84, R^2^ = − 1.30), as did predictions in the opposite direction (negative clip of study 1 to negative clip of study 2: r = 0.299; p = 0.005, MSE = 296.03, R^2^ = 0.061). The model trained on dynFC of subjects watching the neutral clip of study 1 successfully predicted the EE group signature in the negative clip of study 1 (r = 0.334; p = 0.005, MSE = 30.8, R^2^ = −0.389), as did predictions in the opposite direction (negative clip of study 1 to neutral clip of study 1: r = 0.18; p = 0.024, MSE = 14.553, R^2^ = − 45.167). The model trained with the negative clip of study 2 did not predict the neutral clip (r = −0.144; p = 0.999, MSE = 99.432, R^2^ = −314.827), similarly in the opposite direction, theneutral clip of study 1 to negative movie of study 2 (r = −0.028; p = 0.791, MSE = 401.26, R^2^ = −0.272). Taken together, predictions failed only between the neutral clip of study 1 and the negative clip of study 2 but succeeded between the neutral and negative clip of study 1.

The model trained on dynFC patterns to predict the group signature of ED in subjects watching the negative clip of study 2 successfully predicted the group signature of ED in the negative clip of study 1 (r = 0.233, p = 0.005; MSE = 4.497; R^2^ = −0.357), as did predictions in the opposite direction (negative clip of study 1 to negative clip of study 2: r = 0.167, p = 0.005; MSE = 13.673; R^2^ = − 0.128). Conversely, predictions between negative movie clips (study 1 and 2) with the neutral movie clip of study 1 failed (from the negative clip of study 1 to the neutral clip of study 1: r = − 0.025, p = 0.59; MSE = 1.482; R^2^ = − 0.273; from the negative clip of study 2 to the neutral clip of study 1: r = 0.014, p = 0.60; MSE = 5.891; R^2^ = − 4.184; neutral movie clip of study 1 to negative clip of study 1:r = − 0.109, p = 0.96; MSE = 1.344; R^2^ = − 0.324), neutral movie clip of study 1 to negative clip of study 2: r = − 0.009, p = 0.42; MSE = 15.243; R^2^ = − 0.257). Thus these findings suggest that the neural features predicting the group signature response for ED generalize across different narratives and participants for clips inducing high levels of emotional intensity, but not for neutral contexts.

We also examined the top two regions with the highest number of hub-like connections for all movies combined; those with connections positively associated with EE were the angular gyrus (AngG) and the middle temporal gyrus (MTG), while negative connections were observed in the right dorsolateral prefrontal cortex (dlPFC) and right dorsal posterior cingulate cortex (dPCC). For connections positively correlated with ED were the left AngG and the left ventromedial prefrontal cortex (vmPFC), while connections negatively correlated were the right superior lateral occipital cortex (slOC) and the left dPCC.

Next, we sought to understand which canonical brain networks contributed to the group signature responses. To achieve this, we calculated the proportion of significant ROI connections within each network, divided by the total number of possible connections within that network for each movie and EE and ED signature responses separately (Fig. 4A). Across all movies from study 1 and 2, within visual network connections predominated (40%) among those positively correlated with the EE signature response, while only connections within Visual-SAL (14%) negatively correlated with EE. In contrast, for ED, no significant effects were observed across all movies combined. Across the negative clips only from study 1 and 2, EE correlated positively with Visual-DMN (8%) and negatively mainly with Visual-DMN (13%) connections. In contrast, most of the connections correlating positively with ED in negative clips, were found within SAL connections (13%), while negatively correlated mainly with Visual-DMN (7%) connections (see Table 2; results for individual movies are summarized in Table 1).

**Table 1.** Percentage of ROI-pairs within canonical networks connections significantly predicting the group signature responses for EE and ED listed for each clip separately.

| study | Movie-clip | EE |  | ED |  |
| --- | --- | --- | --- | --- | --- |
|  |  | Positive corr. | Negative corr. | Positive corr. | Negative corr. |
| study 1 | Negative clip of study 1 (21 grams) | Visual-Visual (52%) | Visual-SAL (21%) | Visual-Visual (26%) | Visual-SAL (8%) |
|  |  | Visual-DAN (10%) | Visual-DMN (18%) | SMN-SMN (8%) | Visual-CEN (5%) |
|  | Neutral clip of study 1 (Son's room) | Visual-Visual (23%) | Visual-Visual (17%) | Visual-Visual (23%) | Visual-DAN (5%) |
|  |  | Visual-DAN (5%) | SubC-SubC (10%) |  |  |
| study 2 | Sophie's | Visual-Visual (29%) | Visual-SAL (14%) | SAL-SAL (17%) | Visual-DMN (8%) |
|  | choice | DAN-DAN (22%) | Visual-SMN (13%) | SMN-SAL (15%) | Visual-Visual (5%) |

Note: Percentage of ROI-pairs within a canonical network significantly predicting the group signature responses for EE and ED. The two most prominent network pairs are shown.

**Table 2.**
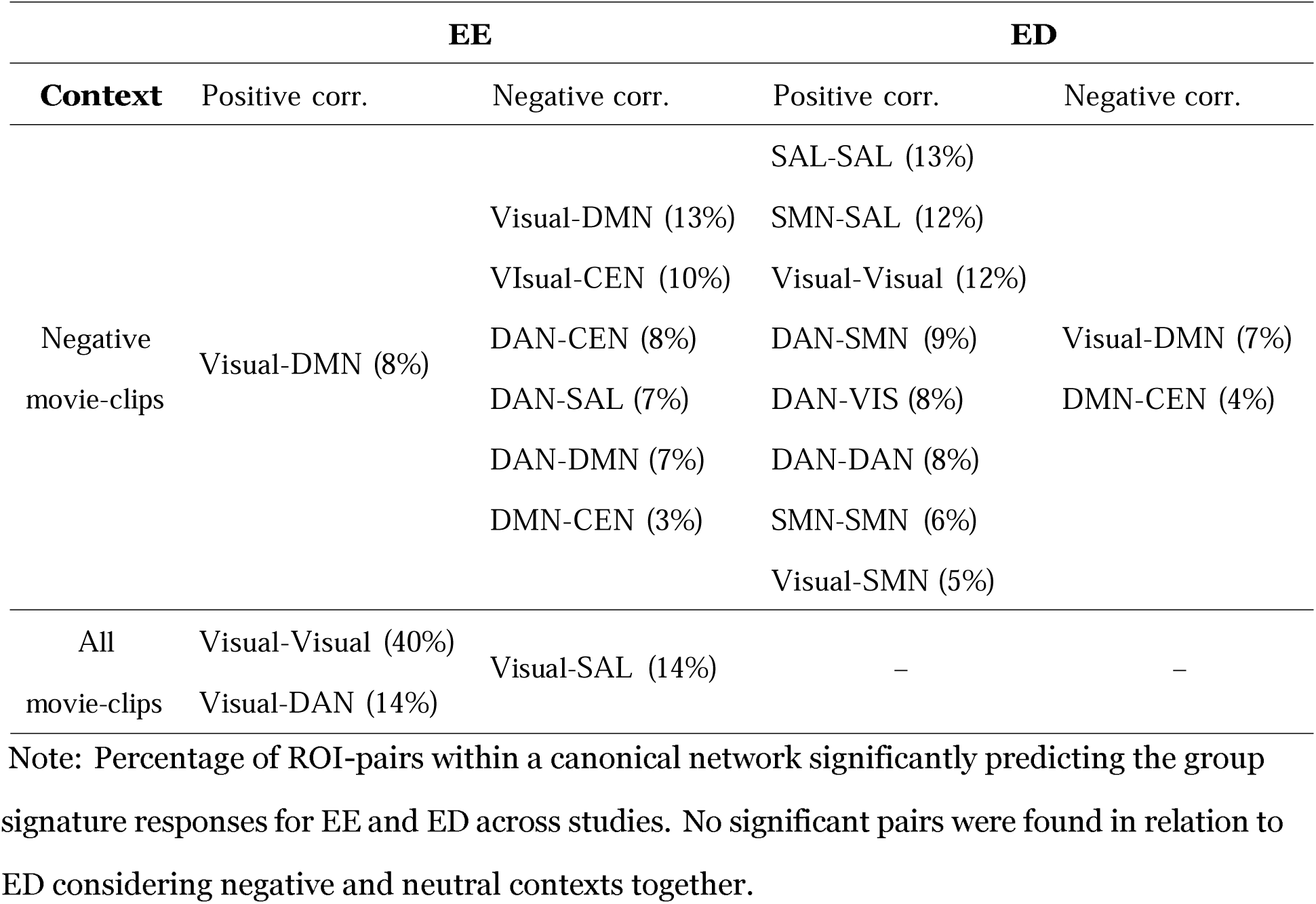
Percentage of ROI-pairs within a canonical network significantly predicting the group signature responses for EE and ED across studies.

## Discussion

This study aimed at identifying generalizable functional connectivity (dynFC) patterns that underlie moment-to-moment changes in collective EE and ED during movie viewing. To do so, we used predictive modeling across movie fMRI data and continuous subjective annotations of affect from two independent studies (Borchardt et al., 2018; Raz et al., 2016). Specifically, we trained a dynFC-based model to predict group signatures of EE and ED in one data set and then tested whether the model could generalize to an independent study with a different clip and participant group and vice versa. We further aimed to test whether predictive dynFC patterns are specific to emotionally intense clips or if they also generalize to a neutral context.

### Disengagement

A model trained to predict an ED group signature based on neural features in one negative movie clip successfully predicted an ED group signature in another negative movie clip, from independent samples with different narratives. However, since this predictive ability did not extend to the neutral clip, this may imply some level of specificity to negative or high emotional intensity contexts. We found that ED was linked to widespread positive associations with between-network connections, specifically for negative movie clips. We hypothesized that ED may reflect an internally directed shift of processing involving the regulation of emotional responses. Thus, the positive association with wide-spread network connections may reflect enhanced cognitive regulation of emotions that relies on the involvement of large scale network communication (Cohen & D’Esposito, 2016; Morawetz et al., 2020). Within these wide-spread connections, the strongest effects were found within the SMN and SAL and between them. Previous research has linked stronger recruitment of the sensorimotor cortex and regions of the salience network to attention directed toward interoceptive markers of arousal (Critchley et al., 2004; Wu et al., 2019). Thus, in our findings, the connections between SAL and SMN during negative emotional movies could indicate an intensified focus on bodily sensations induced by intense emotional experiences. This heightened interoception could play a role in ED, shifting attention away from external emotional stimuli towards internally directed processing and regulation of feeling states. We found that moment-to-moment fluctuations in the ED signature correspond to hub-like behavior within the ventromedial prefrontal cortex (vmPFC). The vmPFC has been implicated in both generating and regulating negative emotions through the formation of generalizable representations of negative emotions (Kragel et al., 2018). Since the highly connected nodes were exclusive to negative movie clips, the involvement of the vmPFC may facilitate the moment-to-moment regulation of negative emotions elicited by negative clips. In our previous study, we observed that inter-individual similarity in static vmPFC activations supported synchronized responses in both EE and ED in the negative clip of study 1 (Nanni-Zepeda et al., 2024). In contrast, adopting a dynamic perspective, the current study reveals that moment-to-moment connectivity of the vmPFC may also reflect fluid adjustments in regulatory processes as emotional contexts shift throughout the movie. Taken together our findings suggest generalizable dynFC patterns across wide-spread large scale networks predicting group collective moment-to-moment changes of ED confined to emotionally intense contexts. However, the specificity of these patterns to negative emotional content may also reflect the particular narrative structures and cinematic techniques employed in the selected clips, which could influence the generalizability across different storytelling approaches or cultural contexts

### Engagement

In contrast to ED, prediction success for EE varied: predictions were successful across negative clips in both study 1 and study 2, and also across the neutral and negative contexts in study 1. However, we interpret the latter finding with caution. A sensitivity analysis varying the width of the thresholds to define EE and ED phases revealed that EE prediction success between neutral and sad movies of study 1 was inconsistent across thresholds (see Table S1). Additionally, predictions did not succeed between the negative clip of study 2 and the neutral clip of study 1.

However, predictions did not succeed between the negative clip of study 2 and the neutral clip of study 1. Thus, idiosyncratic features within the participant group may have driven the successful cross-predictions across clips within one and the same study. Another or complementary interpretation could be that EE relies on network connections that support engagement regardless of emotional intensity levels. However, given that the predictions did not succeed between the neutral clip of study 1 and the negative clip of study 2 the latter interpretation may be more reasonable. Connections within the visual network and between the visual and the DAN were predictive of The EE group signature response.

These networks are critical for goal-directed and stimulus-driven attention, aligning with their established roles in the voluntary control of visuo-spatial attention (Corbetta & Shulman, 2002; Kelley et al., 2008; Tosoni et al., 2023). Specifically, the connectivity in these networks may reflect attentional engagement with the emotional content of emotionally salient stimuli, potentially through mechanisms of visuospatial attention prioritizing emotionally salient stimuli. This prioritization could involve the DAN initiated selective top-down enhancement of visual processing areas linked to the perception and interpretation of emotionally relevant cues (Mohanty & Sussman, 2013). Interestingly, some Visual-to-DMN connections predicted EE in negative and positive directions, highlighting their dual role in promoting and reducing engagement. Prior research has shown that connectivity between the visual cortex and DMN changes depending on task demands, with positive coupling in some contexts like processing task irrelevant information (Chadick & Gazzaley, 2011) or automated information processing (Vatansever et al., 2017), and negative coupling during low cognitive effort (Weber et al., 2022).

Further, during movie viewing, FC profiles of visual-DMN have shown to contribute to the classification of emotional states such as sadness or happiness (Xu et al., 2023). Together these findings may reflect the DMN’s role in facilitating coordination between the DAN and visual networks to direct attention on emotionally salient stimuli.

### Limitations

Using multiple independent datasets introduces variability in cultural background, image acquisition, and annotation methods. The movie clips differed in duration (4.45, 4.54, and 10 min), which may engage distinct neural processes related to sustained attention or narrative structure. However, our analyses target the temporal dynamics of emotional engagement and disengagement (EE/ED) episodes rather than cumulative responses, and the high-intensity scenes driving our measures occurred with similar timing across clips. Different TRs (3s vs. 2s) still yielded comparable numbers of fMRI timepoints, though ideally, clips of equal length would eliminate this confound. Second, EE/ED was derived from retroactive annotations using two methods: continuous pointer ratings and button presses. Both captured the dynamics of interest, but standardized response protocols would improve comparability. While repeated viewings can alter absolute arousal intensity (Chun et al., 2020), we focused on directional changes. Moreover, brain responses show strong test–retest reliability across viewings (Gruskin & Patel, 2022), though it remains unclear whether arousal episodes shift in timing. Third, although our sample was sufficient to detect main effects, larger and more diverse cohorts would improve generalizability and enable study of individual differences. One dataset included only women, and all clips carried a sad affective tone, limiting extension to other populations and emotions. Broader emotional valence should be incorporated in future work. Finally, technical factors may constrain interpretation. Lower-density head coils (8- and 12-channel) could reduce sensitivity to subcortical regions, and stimulus characteristics (narrative complexity, visual features, cultural context) may restrict generalizability to other forms of emotional experience. Future studies with higher-density coils and a wider range of stimuli will provide greater comprehensiveness.

### Conclusion

Our findings reveal that emotional disengagement is characterized by neural patterns specific to emotionally intense situations, while emotional engagement shows broader neural responses that generalize across both high- and low-intensity contexts. This suggests that the neural mechanisms underlying engagement are more adaptable across diverse intensity contexts, whereas disengagement involves more processes specific to high-intensity emotions. These results provide insights into the neural responses driving shifts in emotional experiences and highlight generalizable neural patterns that can guide future research on emotional transitions and regulation in real-world scenarios. Future studies should examine inter-individual variability in model performance to better understand performance heterogeneity across participants and enhance generalizability at the individual level.

## Acknowledgment

We thank our collaborators Simone Grimm from Charité - Universitätsmedizin Berlin and Gal Raz from Tel Aviv University for sharing their data. We used ChatGPT and Perplexity AI for spell checks and help with phrasing and literature search assistance.

## Author contributions

*Melanni Nanni-Zepeda:* Investigation, Data Curation, Formal analysis, Methodology, Software, Writing- Original Draft, review & editing. *Travis C. Evans:* Writing- Reviewing and Editing. *Audreyana Jagger-Rickels:* Writing- Reviewing and Editing. *Gal Raz*: Data source, *Talma Hendler* : Data source, *Yan Fan*: Data source, *Simone Grimm*: Data source, *Martin Walter* : Resources, Supervision. *Michael Esterman:* Writing- Reviewing and Editing. *Agnieszka Zuberer:* Conceptualization, Supervision, Writing- Reviewing and Editing.

## Conflict of interest

The authors declare that there are no conflicts of interest

## Data availability statement

The datasets generated and analyzed during the current study, as well as the custom analysis codes, are available from the corresponding author upon reasonable request. The main analysis scripts were adapted from publicly available code by Song et al. (available at https://github.com/hyssong/NarrativeEngagement).

## Notes

### Competing Interest Statement

The authors have declared no competing interest.

### Summary of Updates

We revised the manuscript to improve clarity and precision in the Methods, Results, and Discussion sections, including rewording and expanded explanations of the analyses and their interpretation. In addition, we added a new supplementary analysis to test the robustness of the main findings using alternative thresholds for defining engagement and disengagement.

