## Supplementary data for "Generalizable Neural Models of Emotional Engagement and Disengagement"

Appendix Supplementary Materials

### Methods

**Instruction for subjective emotional intensity annotations in study 1.**	The instructions were as follows: "Following, are 2 short ratings of the movie clips that you previously watched. You are going to see both movie clips again on the PC screen and are subsequently asked to rate how emotional you felt while watching" Original instruction in German: ’Nun haben wir noch 2 kurze Ratings zu den Filmausschnitten, die sie gerade gesehen haben. Sie werden die Ausschnitte nun nochmal auf dem PC sehen und können dabei mit der Maus eine persönliche Bewertung abgeben, wie stark Sie Momente in dem vorherigen Filmausschnitt bewegt haben.’)

### fMRI Acquisition

In study 1, functional MRI images were acquired on a Siemens Trio 3-T scanner with a twelve-channel radiofrequency head coil, using a T2*-weighted Echo Planar Imaging (EPI) sequence (repetition time (TR)/error time (TE) = 2,000/30 ms, flip angle = 70°, 64× 64 matrix, field of view (FOV) = 192 × 192 mm^2^, in-plane resolution = 3 mm^2^, slice thickness = 3mm, 37 axial slices). The natural-viewing sessions consisted of 147 volumes. T1-weighted anatomical reference images were acquired using 3D-MPRAGE sequence (176 sagittal slices covering the whole brain, 1 mm^3^ isotropic resolution, TR/TE = 1900/2.52 ms, 9° flip angle, 256 × 256 matrix). No headphones were used.

In study 2, functional MRI images were obtained by a GE 3-T Signa Excite echo speed scanner with an eight-channel head coil. Functional whole-brain scans were performed in interleaved order with a T2*-weighted gradient EPI pulse sequence (TR/TE

= 3,000/35 ms, flip angle = 90°, pixel size = 1.56 mm, FOV = 200 × 200 mm, slice thickness = 3 mm, 39 slices per volume). The natural viewing session consisted of 200 volumes. The structural scans included a T1-weighted 3-D axial spoiled gradient echo (SPGR) pulse sequence (TR/TE = 7.92/2.98 ms, 150 slices, slice thickness = 1 mm, flip angle = 15°, pixel size = 1 mm, FOV = 256 × 256 mm). Active noise-cancelling

headphones (Optoacoustic) were used (Raz et al., 2016).

### fMRI Preprocessing

Both data sets were preprocessed using fMRIprep 1.3.2 (Esteban et al., 2019), which is based on Nipype 1.1.9 (K. Gorgolewski et al., 2011; K. J. Gorgolewski et al., 2019). The pipeline included co-registration to the T1w reference using boundary-based registration (Greve and Fischl 2009). Head-motion parameters with respect to the BOLD reference (transformation matrices, and six corresponding rotation and translation parameters) were estimated and regressed out. Spatiotemporal filtering (FSL 5.0.9, Jenkinson et al., 2002) and slice-time corrected using 3dTshift from AFNI 20160207 (Cox & Hyde, 1997). Spatial smoothing with an isotropic, Gaussian kernel of 6mm FWHM (full-width half-maximum) was applied. The BOLD time-series were resampled to MNI152NLin2009cAsym standard space, generating a preprocessed BOLD run in MNI152NLin2009cAsym space. Automatic removal of motion artifacts using independent component analysis (ICA-AROMA) was performed on the preprocessed BOLD on MNI space time-series (Pruim et al., 2015-05-15). Physiological nuisance was corrected by extracting the mean time course of cerebrospinal fluid (CSF) and white matter (WM) calculated from the preprocessed images. The first two volumes of fMRI sequence of movie clips in study 1 were discarded to avoid T1 saturation effect. The last four volumes of fMRI during the negative movie clip in study 1 were removed, given that the clip length was shorter than the entire scan session. One subject from study 1 was discarded due to excessive head movement, identified with mean frame-wise displacement (threshold = 0.20) resulting in 21 participants. Applying the same cut-off to study 2, three subjects were discarded resulting in forty-four participants in the final sample.

**Robustness of Engagement and Disengagement Predictions to Varying Percentile Thresholds.**	To assess the robustness of our engagement and disengagement predictions, we tested whether the results held across varying percentile thresholds used to

define the phases. The original threshold range (low: 33rd percentile, high: 67th percentile) was modified to include both wider and narrower ranges. Specifically, we expanded the range in three steps by lowering the low percentile and increasing the high percentile in increments/decrements of 5 units, resulting in the following additional threshold pairs: Percentile 1 (low: 28, high: 72), percentile 2 (low: 23, high: 77), and percentile 3 (low: 18, high: 82). To test the opposite direction, we also applied a narrower range with percentile 4 (low: 38, high: 62). However, additional narrowing resulted in excessive data loss and was not pursued further. Examples of engagement and disengagement estimation using these alternate thresholds are illustrated in Figure S3 for selected participants, highlighting comparisons to the original range.

**Results:** The full results from prediction analyses using each percentile pair are summarized in Table S1 for EE and Table S2 for ED. For engagement, predictions between the two sad movies (21 Grams and Sophie) remained robust across all four threshold ranges. Regression models trained on one sad movie and tested on the other consistently yielded significant results, replicating the pattern observed with the original percentile thresholds. This supports our primary finding that engagement during sad movies exhibits a stable and generalizable temporal structure. In contrast, predictions between sad and neutral movies showed more variability. Significant correlations emerged in some, but not all threshold ranges. Specifically, models trained on 21 Grams and tested on the neutral movie yielded significant results for percentiles 2 and 3. The reverse prediction (neutral to 21 Grams) showed significance only for percentile range 2. No significant predictive relationship was found between Sophie and the neutral movie in either direction, regardless of the threshold used. Taken together, as in our original findings, cross-movie predictions presented strong generalization only between sad movies, while predictions across movies with differing emotional valence (sad vs. neutral) were less consistent and more sensitive to threshold definitions.

Disengagement predictions were robust across all four threshold ranges for the

sad-sad movie comparisons, replicating the original findings. Similar to the engagement results, no significant correlations were observed for predictions between sad and neutral movies, further confirming the specificity of disengagement dynamics within the same emotional context.

**Conclusion:** Overall, these control analyses confirm the robustness of our main findings to variations in the engagement/disengagement definition. The only variability observed was in engagement predictions between the sad movie from study 1 (21 Grams) and the neutral movie, where not all percentile thresholds yielded significant results. This reinforces our original hypothesis that temporal engagement patterns are only reliably predictable between sad movies across independent studies

### Results

**Distinctive Patterns of Emotional Engagement and Disengagement.**	We expected that the negative clips would induce more emotional shifts (EE and ED) than the neutral clip. To test this, we built linear mixed-effects model to test if the number of shifts could be predicted by the interaction between movie clip and shift direction (EE and ED). This interaction was significant (*β* = 1.78; 95% CI: [1.19, 2.37], *p < .*001). A post-hoc test showed that the effect was significant in the negative movie of study 1 (estimate[ED-EE]=

-2.41 ∓0.429; 95% CI: [-3.26, -1.51], *p < .*001), negative movie of study 2

(estimate[ED-EE]= -1.78 ∓0.300; 95% CI: [-2,37, -1.18], *p < .*001), but not significant for

the neutral movie of study 1 (estimate[ED-EE]= -0.50 ∓0.429; 95% CI: [-1.35, 0.348],

*p* = *.*245) (see Fig. S1). We can conclude that (1) the number of emotional shifts was higher during the negative compared to neutral condition for EE and ED and (2) the number of EE was higher than ED in negative condition with no significant difference in the neutral condition.

In the negative clip of study 1, participants reported an average of 6.90∓4.82 EE time points (4.9% ∓3.4%), and 2.40∓3.14 ED time points (1.7% ∓ 2.22%), with a mean

duration of 13.81∓9.65 seconds for EE and 4.81∓6.28 seconds for ED. For the negative clip of study 2, participants reported an average of 3.8∓2.46 EE time points (1.9% ∓1.23% ) and 1.4∓1.42 ED time points (0.7% ∓ 0.71%), with a mean duration of 11.4∓7.39 seconds for EE and 4.2∓4.26 seconds for ED. In the neutral clip of study 1, there were, on average,

1.31∓1.64 EE time points (0.9%∓1.12%) and 0.9∓1.63 ED phases (0.62%∓1.11%), with a mean duration of 2.63∓3.28 seconds for EE and 1.81∓3.26 seconds for ED. In each movie clip, some participants did not exhibit EE time points (3 subjects from the negative movie

clip of study 1 and 10 subjects from the neutral movie of study 1), and some did not have ED time points (10 subjects from the negative movie clip of study 1, 15 subjects from the negative movie clip of study 2, and 13 subjects from the neutral movie clip of study 2).

We expected that emotional movies would induce more emotional shifts (EE and ED) than the neutral clip. To test this, we built linear mixed-effects models by choosing a

step-up approach (Hox et al., 2017) including an effect in the model if it significantly improved the model fit. A model fit was improved if the difference between the

log-likelihood ratio of a model that contained the effect and a model that did not, was significant (as compared with ANOVA; p<0.05). We used mixed effects modeling to test if the number of shifts could be predicted by emotional condition (negative/neutral) and emotional shift (EE/ED) with subjects as a random intercept, and movie duration as a fixed effect. The results revealed a significant interaction effect between emotional context and emotional shift (*β* = 1.18; 95% CI: [0.17, 2.18], *p* = *.*02). A post-hoc test revealed that

(1) the number of emotional shifts was higher during the negative condition compared to

neutral condition for both EE and ED (EE: estimate[neutral-negative]= -2.60; ∓0.411 95% CI: [-3.41, -1.79], *p < .*001; ED: estimate [neutral-negative]= -1.42; ∓0.486 95% CI: [-2.39,

-0.46], *p* = *.*004); and (2) the number of EE was higher than ED in negative condition

(estimate[ED -EE] = -1.68; ∓0.275; 95% CI: [-2.22, -1.13], *p < .*001) with no significant difference between EE and ED in the neutral condition (estimate[ED-EE]= -0.50; ∓0.428 95% CI: [-1.35, 0.34], *p* = *.*244)

**Within-dataset predictions.**	SVR models successfully predicted the corresponding group signature responses for both EE and ED within all three movie clips at levels significantly above chance. For EE, the strongest association was observed for the negative movie in study 1 . (*r* = 0*.*823*, p <* 0*.*001; *MSE* = 0*.*454*, R*^2^ = 0*.*546), followed by the negative movie in study 2 (*r* = 0*.*722*, p <* 0*.*001; *MSE* = 0*.*552; *R*^2^ = 0*.*448), and the neutral movie of study 1 (*r* = 0*.*618*, p <* 0*.*005; *MSE* = 0*.*663*, R*^2^ = 0*.*337). Similarly, for ED, , significant above-chance predictions were achieved within all conditions: negative movie study 1 (*r* = 0*.*689*, p <* 0*.*001; *MSE* = 2*.*673; *R*^2^ = 0*.*331), negative movie of study 2 (*r* = 0*.*593*, p <* 0*.*001; *MSE* = 8*.*444; *R*^2^ = 0*.*302) and the neutral movie of study 1

(*r* = 0*.*516*, p <* 0*.*005; *MSE* = 0*.*778*, R*^2^ = 0*.*222) see Fig. S2.

*Table S1*

*Emotional Engagement*

**Percentiles range**

Percentile 1

low: 28, high: 72

Percentile 2

low: 23, high: 77

Percentile 3

low: 18, high: 82

Percentile 4

low: 32, high: 62

**Dataset learn: Sophie Dataset test: Grams** pearson r = 0.385** MSE = 36.061

r-squared = 0.084 pearson r = 0.444** MSE = 38.895

r-squared = 0.135 pearson r = 0.522** MSE = 29.875

r-squared = 0.161 pearson r = 0.604** MSE = 6.008

r-squared = -0.225

**Dataset learn: 21 Grams Dataset test: Sophie** pearson r = 0.167**

MSE = 107.124

r-squared = 0.017 pearson r = 0.228** MSE = 93.676

r-squared = 0.042 pearson r = 0.266** MSE = 86.922

r-squared = 0.054 pearson r = 0.303** MSE = 41.155

r-squared = 0.0

**Dataset learn: 21 Grams Dataset test: Sophie**

**Dataset learn: 21 Grams Dataset test: Neutral** pearson r = 0.103

MSE = 29.282

r-squared = -8.764 pearson r = 0.179* MSE = 23.73

r-squared = -8.66 pearson r = 0.190* MSE = 18.81

r-squared = -16.292 pearson r = 0.112 MSE = 1.619

r-squared = -3.094

**Dataset learn: Neutral Dataset test: 21 Grams** pearson r = 0.059

MSE = 60.977

r-squared = -0.539 pearson r = 0.154* MSE = 61.454

r-squared = -0.358 pearson r = 0.087 MSE = 48.94

r-squared = -0.365 pearson r = -0.075 MSE = 5.644

r-squared = -0.119

**Dataset learn: Neutral Dataset test: Sophie** pearson r = -0.191

MSE = 142.129

r-squared = -0.304 pearson r = -0.131 MSE = 118.847

r-squared = -0.215 pearson r = -0.11 MSE = 110.472

r-squared = -0.202 pearson r = -0.212 MSE = 47.087

r-squared = -0.144

**Dataset learn: Sophie Dataset test: Neutral** pearson r = -0.131

MSE = 41.102

r-squared = -12.729 pearson r = -0.098 MSE = 27.323

r-squared = -10.168 pearson r = -0.06 MSE = 28.927

r-squared = -25.664 pearson r = -0.037 MSE = 8.423

r-squared = -20.338

* *p <* 0*.*05, ** *p <* 0*.*01

*Table S2*

*Emotional Disengagement*

**Percentiles range**

Percentile 1

low: 28, high: 72

Percentile 2

low: 23, high: 77

Percentile 3

low: 18, high: 82

Percentile 4

low: 32, high: 62

**Dataset learn: Sophie Dataset test: 21 Grams** pearson r = 0.383**

MSE = 2.108

r-squared = 0.067 pearson r = 0.3** MSE = 2.128

r-squared = -0.022 pearson r = 0.268** MSE = 1.686

r-squared = -0.05 pearson r = 0.182** MSE = 1.093

r-squared = -2.718

**Dataset learn: 21 Grams Dataset test: Sophie** pearson r = 0.279**

MSE = 7.783

r-squared = 0.059 pearson r = 0.286** MSE = 5.398

r-squared = 0.065 pearson r = 0.304** MSE = 3.779

r-squared = 0.069 pearson r = 0.281** MSE = 1.946

r-squared = -0.146

**Dataset learn: 21 Grams Dataset test: Sophie**

**Dataset learn: 21 Grams Dataset test: Neutral** pearson r = -0.037

MSE = 8.106

r-squared = -0.183 pearson r = -0.027 MSE = 3.958

r-squared = -0.116 pearson r = -0.002 MSE = 2.589

r-squared = -0.067 pearson r = -0.052 MSE = 0.719

r-squared = -0.262

**Dataset learn: Neutral Dataset test: 21 Grams** pearson r = -0.092

MSE = 3.99

r-squared = -0.778 pearson r = -0.122 MSE = 2.719

r-squared = -0.323 pearson r = -0.095 MSE = 1.857

r-squared = -0.169 pearson r = -0.008 MSE = 0.529

r-squared = -0.796

**Dataset learn: Neutral Dataset test: Sophie** pearson r = 0.003

MSE = 9.755

r-squared = -0.18 pearson r = -0.071 MSE = 6.445

r-squared = -0.117 pearson r = -0.09 MSE = 4.383

r-squared = -0.08 pearson r = -0.1 MSE = 1.801

r-squared = -0.06

**Dataset learn: Sophie Dataset test: Neutral** pearson r = 0.03

MSE = 8.129

r-squared = -0.187 pearson r = -0.031 MSE = 4.217

r-squared = -0.195 pearson r = -0.039 MSE = 2.792

r-squared = -0.165 pearson r = -0.094 MSE = 0.981

r-squared = -0.765

* *p <* 0*.*05, ** *p <* 0*.*01

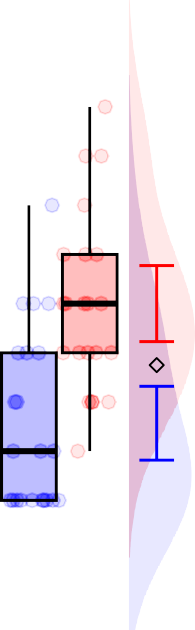

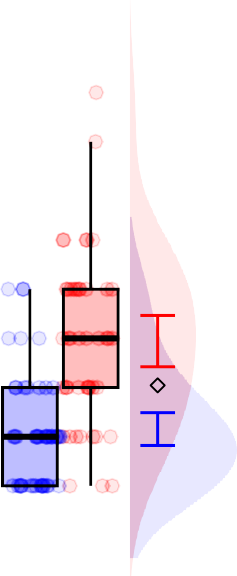

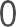

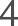

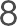

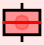

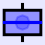

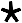

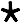

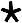

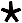

EE

ED

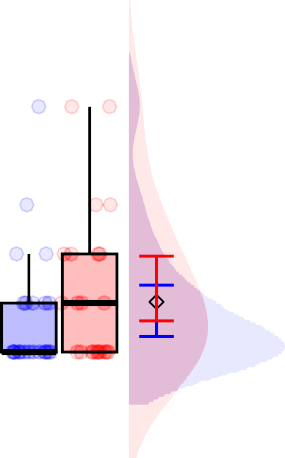
21 grams (negative)

Number of period blocks

Sophie's choice (negative)

Son's room (neutral)

*Figure S1.* Estimated number of emotional engagement (EE) and emotional disengagement (ED) periods, inferred from fluctuations in subjective emotional intensity annotations across the three movie clips

**Engagement**

**Within movie prediction**

**Disengagement**

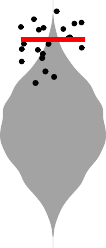

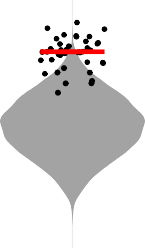

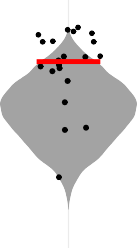

1.0

r (predicted, observed)

0.5

0.0

-0.5

r= 0.823

p= 0.001

r= 0.722 p=0.001

r= 0.618 p=0.005

0.5

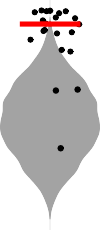

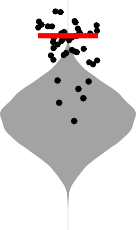

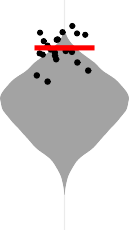

0.0

-0.5

r= 0.689 p=0.001

r= 0.593

p= 0.001

r= 0.516

p= 0.005

Grams	Sophie	Son's room	Grams	Sophie	Son's room

*Figure S2.* Model performance for within-dataset validations. The black dots indicate Pearson’s correlations between predicted and observed engagement (top) or disengagement (bottom) for every participant. Horizontal lines indicate the quality of prediction as reflected in the mean r across cross-validation folds. Violin plots depict the null distributions of mean prediction performance of models predicting engagement and disengagement time courses after

phase-randomization. The significance of empirical r was computed based on the null distribution (one-tailed t tests).
